# Correlates of time to first citation in ecology and taxonomy

**DOI:** 10.1101/2023.03.16.532892

**Authors:** Jhonny J. M. Guedes, Isabella Melo, Igor Bione, Matheus Nunes

## Abstract

Several metrics exists to evaluate the impact of publications and researchers, but most are based on citation counts, which usually fail to capture the temporal aspect of citations. Time to first citation represents a useful metric for research evaluation, and informs the speed at which scientific knowledge is disseminated through the scientific community. Understanding which factors affect such metrics is important as they impact resource allocation and career progression, besides influencing knowledge promotion across disciplines. Many ecological works rely on species identity, which is the ‘coin’ of taxonomy. Despite its importance, taxonomy is a discipline in crisis lacking staff, funds and prestige, which ultimately may affect the evaluation and dissemination of taxonomic works. We used a time-to-event analysis to investigate whether taxonomic, socioeconomic, and scientometric factors influence first citation speed across hundreds of ecological and taxonomic articles. Time to first citation differed greatly between these areas. Ecological studies were first cited much faster than taxonomic studies. Multitaxa articles received first citations earlier than studies focused on single major taxonomic groups. Article length and *h*-index among authors were negatively correlated with time to first citation, while the number of authors, number of countries, and Gross Domestic Product was unimportant. Knowledge dissemination is faster for lengthy, multitaxa, and ecological articles relative to their respective counterparts, as well as for articles with highly prolific authors. We stress that using several unrelated metrics is desirable when evaluating research from different–and even related–disciplines, particularly in the context of professional progression and grant allocation.

## Introduction

The publication of scientific research is an essential step for the progress of science because it allows for discussions within the scientific community about what has been done in a particular field. The number of scientific articles published annually, both globally and in specific areas, grows at high rates (Bornmann & Mutz 2015; Larsen & von Ins 2010). However, the high volume of publications and, therefore, of research being performed, implies that limited financial resources for scientific research (especially in developing countries) may become increasingly scarce—i.e., if the increase of research projects is not followed by a similar increase in research funding. Hence, resource allocation usually follows some kind of technical criteria. Indeed, several metrics—mainly based on the number of citations (Aksnes et al. 2019; Garfield 1972)—were developed in an attempt to assess academic productivity and the impact of scientific research. Despite numerous criticisms about the inadequacy of these metrics in evaluating the impact of scientific output and researchers in some areas [e.g., medical sciences (Dinis-Oliveira 2019), taxonomy (Pyke 2014; Valdecasas et al. 2000), social sciences and humanities (Steele et al. 2006)] as well as their impact outside academia (Ravenscroft et al. 2017), these metrics are still commonly used by research funding institutions to decide where resources will be allocated (Fortin & Currie 2013).

Several metrics have been proposed throughout the years and they may ‘behave’ differently between areas as well as be affected by different factors (Padial et al. 2010; Tahamtan et al. 2016). In ecology, for example, variables such as number of authors, journal impact factor, article size (number of pages), study results (positive or negative outcome), among others, may influence the total number of citations (Borsuk et al. 2014; Leimu & Koricheva 2005). However, some variables may be good predictors of the number of citations in some areas, but not in others, so features of the study area in which the work is inserted and its respective ‘audience’ represents another important factor to be considered (Aksnes et al. 2019; Krell 2002). For instance, taxonomy differs from other areas of the biological sciences due to several factors, such as the decrease in professional taxonomists over time (Hopkins & Freckleton 2002; Wägele et al. 2011) and the increase in the number of specialists in particular taxonomic groups (Joppa et al. 2011), with little interaction between experts working in different groups (Venu & Sanjappa 2011). These factors may reduce the short-term citation potential of taxonomic articles compared to articles from other disciplines. That is, the process of citation accumulation in taxonomy is usually slow and gradual, and may take decades to reach levels quickly observed in other disciplines (Venu & Sanjappa 2011). Therefore, interdisciplinary differences in citation patterns must be taken into account during, for example, resource allocation and professional progression.

Understanding which factors influence scientometric indices is important because only then will we know under which contexts and areas these metrics will be relevant. However, most of these metrics reflect ‘snapshots’ that do not characterize the temporal behaviour of citations (Schubert & Glanzel 1986). For example, some articles—called ‘sleeping beauty’—receive belated recognition through citations by the academic community, while others are immediately noticed, receive many citations, but are not much cited in the following years (Van Dalen & Henkens 2005). Thus, it is also important to understand the patterns and processes underlying the reception speed and recognition—or level of acceptance—of articles by the scientific community, which can be measured by the time to first citation (Bornmann & Daniel 2010a; Van Dalen & Henkens 2005; Egghe et al. 2011; Glanzel & Rousseau 2012; Kumari et al. 2020; Schubert & Glanzel 1986). In this sense, both the impact—thorough citation counts—and the time to first citation are important metrics to consider when analysing the scientometric aspects of publications (Van Dalen & Henkens 2005). Still, the later could be, for instance, an early indicator of scientific performance (Kumari et al. 2020). Although we have a good understanding of which factors affect the total number of citations —and related metrics— in numerous areas, studies analysing which factors affect the time to first citation and how it correlates with total citations are still scarce (Hancock 2015; Kumari et al. 2020). So far, we know that in some disciplines such times decrease with an increase in the number of authors, differs drastically between research areas, and increases if the article was rejected before being published (Bornmann & Daniel 2010a; Glanzel & Rousseau 2012; Hancock 2015).

Herein, we aim to understand which factors influence the time to first citation of articles published in the areas of ecology and taxonomy. These two related—but distinct— areas within biology interact directly with each other once, for instance, many ecological analyses rely on species identity, such as regional species lists (Nekola & Horsák 2022), which in turn is the ‘coin’ of taxonomy. Specifically, we will test the potential effects of the i) number of pages, ii) number of authors, iii) number of countries (according to authors’ addresses), iv) maximum *h*-index among authors (this index measures both productivity and impact, in terms of citations, from the publications of a researcher), v) maximum per capita gross domestic product (GDP) of affiliation countries among authors, vi) study area, and vii) taxonomic group studied. We expect the time to first citation will decrease with an increase in the number of pages (greater amount of information presented), the number of authors and countries (higher collaboration or multidisciplinarity), *h*-index (higher impact authors), and GDP (greater influence of developed countries). Moreover, we expect lower times to first citation in ecological than taxonomic articles due to intrinsic differences between areas as mentioned above. We expect that studies on more charismatic groups, such as animals and plants, which have more researchers working on them and are more appealing to the public (Troudet et al. 2017), will receive their first citation faster than more neglected groups, such as microorganisms (e.g. bacteria, protists). Lastly, we also discuss how this metric correlates to the total number of citations among studies. Overall, this work seeks to improve our understanding of potential factors influencing one of the many scientometric aspects of scientific publications: the speed of the first citation.

## Methodology

### Data collection

To investigate potential factors influencing the time to first citation in taxonomic and ecological works, we chose ten journals from each area (n = 20 journals; see Appendix 1) and randomly selected using the *sample_n* function, without replacement, from the *dplyr* package in software R (Wickham et al. 2020), 30 publications from the year 2015 for each journal (*n* = 600 articles). We excluded articles that did not fit the study goals—e.g., theoretical or simulation articles that did not analyse any species—and replaced them (also using the R *sample_n* function) with other articles from the same journal to keep 30 publications per journal. We selected journals based on three criteria: i) journals that were important channels of information diffusion in each area; ii) journals whose scope involved only one of the areas of interest, that is, we avoided multidisciplinary journals because it is difficult distinguishing whether a paper was taxonomic or ecologic in scope, and iii) journals that were indexed in the Web of Science (hereafter, WoS) database. In the case of taxonomy, we also selected journals known for publishing many articles on species descriptions or natural history.

We searched the publications of each journal from the year 2015 through the WoS database (search performed on 24 April 2021). Given we are interested in the time to first citation, the six-year time period is sufficiently long to allow us to study the phenomenon of interest. We used publications of the same year to reduce variation in potential correlates of time to first citation, and to allow more direct comparisons among publications analysed herein. Our search in WoS returned the publication list and several scientometric data related to each publication, some of which were later used as predictors in our analysis (see below). All publications represented either original articles or revisions (based on the ‘document type’ category provided by WoS). For each one, we compiled data about eight potential predictors of the time to first citation. Six were continuous covariates: 1) number of pages; 2) number of authors; 3) number of countries (based on authors’ addresses); 4) maximum *h*-index among authors (obtained from WoS); 5) maximum per capita GDP (in US$) of affiliation countries among authors—obtained from the World Bank through the R package *wbstats* (Piburn 2020). Two covariates were categorical and informed: 7) the study area (ecology or taxonomy, based on the journal in which the article was published); and 8) the focal taxonomic group studied: plants, animals, multitaxa (i.e., analysing species from different kingdoms or domains), or ‘others’ (i.e., bacteria, fungi, protists, or chromists). We measured the time to first citation as the number of months between the WoS publication date and the date of first citation (excluding self-citations and considering WoS citation date). Articles that were not cited or only received self-citations by April 2021 (cut-off date) received maximum time to first citation—given their publication and search dates. Lastly, we created a binary variable called ‘censor’ that informed whether articles were cited (1) or not (0) by the cut-off date.

### Statistical analyses

Before performing the statistical analysis, we log_10_-transformed some continuous variables (number of pages, number of authors, number of countries, and *h*-index) due to high skewness and kurtosis to increase linearity with our response variable —values between −2 and +2 are considered good evidence of a normal distribution, and hence, do not need transformation (George & Mallery 2010). We standardized all continuous variables (mean = 0, std. deviation = 1) to directly compare their effect sizes and checked for multicollinearity using the Variation Inflation Factors - VIFs (Mansfield & Helms 1982). VIF values higher than 10 indicate that variables should be removed from the analysis (Kutner et al. 2005), but since none of our variables reached VIFs higher than two (Supplementary Table S1), we kept them all.

The time to first citation was investigated through time-to-event analysis, also known as survival analysis (Bradburn et al. 2003; Clark et al. 2003). This analysis is commonly used to investigate factors that may influence the probability of a particular event, such as time to death or recovery, or failure time of equipment. Our event of interest—or survival time—is the time to first citation after publication. Survival analysis incorporates censured data (i.e., observations that lack the event of interest at the end of data collection), which is an advantage from commonly used statistical methods or previously proposed first-citation-speed indexes (Bornmann & Daniel 2010a; Egghe et al. 2011), where censured data is discarded despite containing valuable information. Specifically, we used an Accelerated Failure Time (AFT) survival model, a parametric model that allows the evaluation of covariate effects upon survival times (Bagdonavicius & Nikulin 2002). We applied an AICc-based model selection approach to identify the best family error distribution (exponential, Weibull, lognormal, log-logistic, gamma, and Gompertz) for the AFT model (Burnham & Anderson 2002). The best family error distribution identified was the log-logistic (Supplementary Table S2, Figure S1). Lastly, we checked the relationship between the time to first citation and total citation counts through a linear model to investigate how one may influence another. All analyses and plots were performed/created in the software R 4.0.2 (R Core Team 2020) using the packages *survival* (Therneau 2021), *flexsurv* (Jackson 2016), *survminer* (Kassambara et al. 2021), and *ggplot2* (Wickham 2016).

## Results

Time to first citation across ecological and taxonomic works combined ranged from zero to 75 months, with a median of 14 months. Among all articles analysed, 524 (or 87.3%) were cited at least once by the cut-off date of this study. About one-quarter of all articles received their first citation up to seven months after publication, but another quarter was first cited only more than 30 months (two and a half years) after being published. However, time to first citation varied significantly between study areas (Figure 1a; Table 1). Median survival time across ecological studies was 10 months (range = 0—74), while in taxonomic studies it more than doubled (26 months, range = 0—75). Five years after publication, more than one-quarter of taxonomic works had not been cited once, while less than three percent of ecological works remained to be first cited (Figure 1b).

**Table 1:** Accelerated Failure Time (AFT) survival model of the time to first citation as a function of taxonomic, socioeconomic, and scientometric predictors. Continuous predictors were z-transformed before analysis to make them comparable. Categorical predictors have a baseline level used for comparisons. Taxonomic studies are being compared to ecological ones. Studies with animals are being compared to studies with plants, ‘others’ and multitaxa.

| Predictors | Estimate | Std. Error | z-value | p-value |
| --- | --- | --- | --- | --- |
| **(Intercept)** | **2.525** | **0.079** | **32** | **<0.001** |
| **LogN.Pages.z** | **-0.170** | **0.049** | **-3.5** | **<0.001** |
| LogN.Authors.z | -0.041 | 0.053 | -0.78 | 0.438 |
| **LogH.index.Max.z** | **-0.203** | **0.060** | **-3.4** | **0.001** |
| LogN.Countries.z | -0.037 | 0.049 | -0.76 | 0.449 |
| GDP.Max.z | 0.009 | 0.052 | 0.17 | 0.868 |
| **Taxonomy** | **0.698** | **0.120** | **5.8** | **<0.001** |
| TaxOthers | 0.129 | 0.171 | 0.76 | 0.449 |
| TaxMultitaxa | -0.159 | 0.131 | -1.21 | 0.225 |
| TaxPlantae | -0.063 | 0.099 | -0.64 | 0.520 |

**Figure 1:**
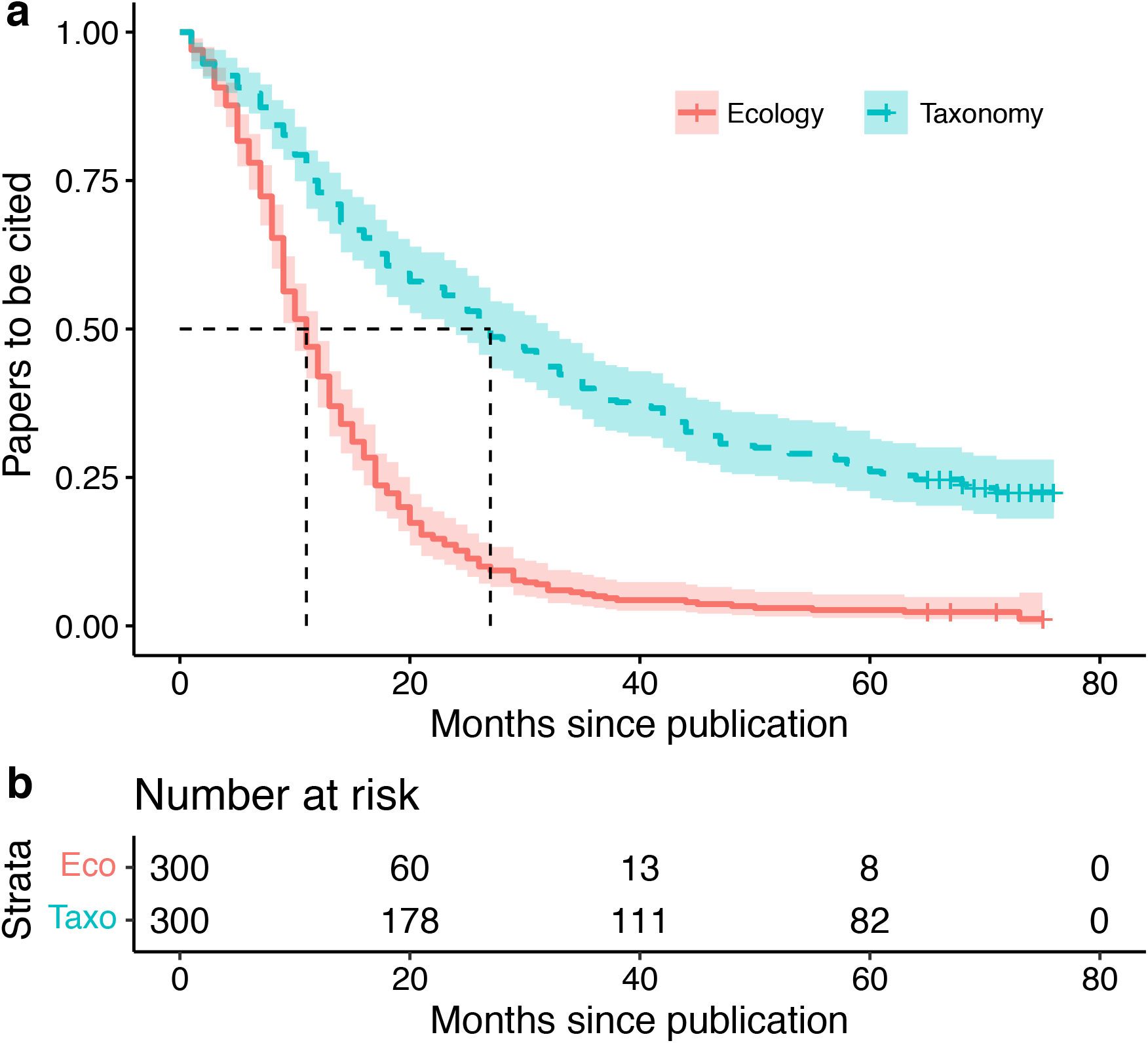
Proportion of ecological and taxonomic articles to receive their first citation after publication. a) Survival curves show probabilities of receiving first citation and their 95% confidence intervals. This probability represents the chance of an article receiving its first citation at time *t*+1 given that it was not cited at time *t*. Black dashed lines indicate the median time to first citation for each study area when survival probability was 50%. b) The number of articles remaining to be cited in *n* months since publication.

We found that multitaxa studies—e.g., studies on interaction networks, food webs— received their first citation much faster than studies focused on species from a single major taxonomic group (Figure 2a-b). Median time to first citation was 9.5 months for multitaxa studies, 14 months for articles focused on animals or plants, and more than two years (28 months) for studies on more neglected groups —mostly microorganisms. However, differences among groups were not significant, likely due to high variability and overlap in survival times (Table 1; Figure 2a).

**Figure 2:**
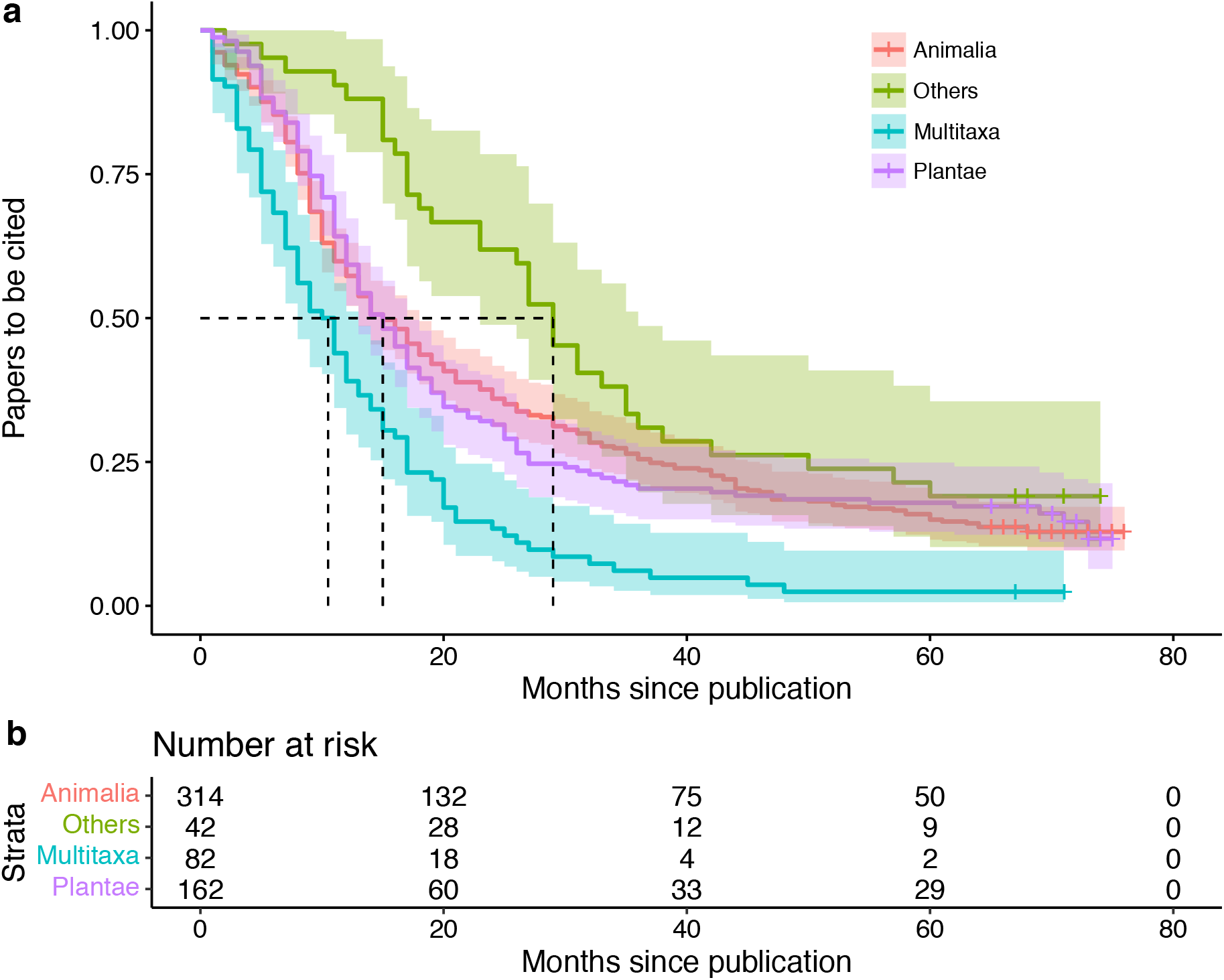
Proportion of articles based on different major taxonomic groups to receive first citation after publication. a) Survival curves show probabilities —of receiving first citation— and their 95% confidence intervals. Details are the same as for figure 1. b) The number of articles remaining to be cited in *n* months since publication. Legend: Others = Bacteria, Chromista, Fungi, and Protista combined.

Aside from the study area, the most important predictor—i.e., strongest effect size— in explaining variation in time to first citation was the maximum *h*-index among authors. Precisely, time to first citation decreases with increasing productivity and impact of researchers. Likewise, the number of pages is negatively associated with the time to first citation. That is, lengthy articles receive, on average, their first citation faster than shorter ones. The number of authors, number of countries, and maximum GDP were unimportant in explaining variation in the time to first citation (Table 1). Overall, our AFT model explained 26.4% of the variation in the time to first citation across ecological and taxonomic articles. Lastly, time to first citation was negatively correlated with log-transformed total number of citations (lm: coef. = −0.032, t = −20.4, p < 0.001), with a coefficient of determination of 41% (Supplementary Figure S2).

## Discussion

We analysed the effects of taxonomic, socioeconomic, scientometric, and field-related predictors on first citation speed across hundreds of ecological and taxonomic articles more than five years since their publication. We have shown that recognition and speed of knowledge dissemination in the scientific community is much faster for articles in ecology than taxonomy. In other words, there is a field-related bias regarding this particular metric. We also showed that first citation speed is, on average, faster for lengthy articles and for papers authored by high-impact researchers, while socioeconomic and collaboration predictors were unimportant. Time to first citation can differ greatly among studies focused on different taxonomic groups, particularly between multitaxa studies and those focused on less charismatic and conspicuous taxa. Moreover, time to first citation shows a negative non-linear relationship with the total number of citations and high variability among articles with intermediate citation counts, which may hamper the use of this metric alone as a good surrogate for future number of citations. Overall, our findings show that time to first citation is lower for lengthy, multitaxa, and ecological articles relative to their respective counterparts. Likewise, papers authored by more productive and prolific authors are likely to receive their first citation earlier.

Since the proposal of using the number of citations as a metric for research assessment (Garfield 1972), several studies have investigated factors that may affect this metric and under which circumstances [for a review, see (Tahamtan et al. 2016)]. Conversely, only a few studies have analysed which factors influence the time to first citation and how it varies among disciplines (Hancock 2015; Kumari et al. 2020). Differently from the total number of citations—usually perceived as the impact of scientific publications (Van Dalen & Henkens 2005)—the time to first citation can be seem as the speed to which knowledge and ideas in a given area are disseminated through the scientific community (Glanzel & Rousseau 2012; Hancock 2015). Thus, although these metrics are somehow related (Supplementary Figure S2) and could, perhaps and to some extent, be used to predict one another (Kumari et al. 2020), they provide information on different patterns about scientific publications. While the number of citations can accumulate through time, usually at distinct rates among disciplines, the time to first citation represents a particular event that somehow levels all publications under comparison based on a specific temporal aspect of citations.

We have shown empirically that taxonomic articles usually receive their first citation much later than ecological papers, which is in line with previous studies showing differences in citation speed between distant and even closely related disciplines (Abramo et al. 2011; Glanzel & Rousseau 2012; Hancock 2015; Oermann et al. 2010). Coupled with the slow and gradual accumulation of citations in taxonomic studies through time (Venu & Sanjappa 2011), our findings highlight the inadequacy of simply using citation counts as a tool for assessing impact and then comparing taxonomic and non-taxonomic studies. If comparisons are needed such as for decisions on grant allocation or career development, alternative and more appropriate metrics —citation-based or not— could be used (Pyke 2014; Valdecasas et al. 2000). For instance, the Field-Weighted Citation Index (Zanotto & Carvalho 2021) is a recently developed metric that compares related articles—i.e., from the same field—based on their keywords and age. Differences in citation counts and time to first citation are expected among areas due to intrinsic differences between them (Aksnes et al. 2019; Krell 2002). For example, community size, publication rates, time for obsolescence, and citation practices adopted in different areas can impact citation counts and other citation-based metrics. Across all selected journals in this study, journal impact factors were much higher in ecology than taxonomy (Supplementary Figure S3a), which highlights yet another difference between areas that can, in turn, also affect time to first citation (Supplementary Figure S3b). Therefore, although using simple metrics is tempting, applying less biased, more comparable, or even a combination of metrics is desirable and should be encouraged, particularly in funding agencies and universities, to reduce existing gaps between disciplines.

We found a negative relationship between time to first citation and maximum *h*-index among authors. The *h*-index was a metric proposed to quantify author-level productivity and impact based on a combination of the number of publications and citations (Hirsch 2005). There are several critics about using this metric as researchers’ scientific achievements for professional progression or funding allocation because, for instance, it does not accommodate age disparities among researchers (Dinis-Oliveira 2019). Although this metric may indeed be age-biased, higher *h*-indexes are likely related to very prolific researchers with equally higher networks within the scientific community that, in turn, seems to contribute to the fast dissemination of produced knowledge and ideas in their scientific field. One point of concern that arises with such finding is that, for instance, good scientific work can be developed by less prolific authors, but such works may take more time—if ever—to gain acknowledgment by their peers. Additionally, high-impact authors may be ‘invited’ to contribute in works where they have actually not contributed at all, be it intellectually or otherwise. Although the latter possibility may be hard to investigate, it warrants attention since such practice can be detrimental to science as a whole.

Similarly to the *h*-index, an increasing number of pages may decrease first citation speed. That is, lengthy articles usually receive their first citation faster than do shorter ones. Articles’ length is also known to affect citation frequency in ecology (Leimu & Koricheva 2005; Padial et al. 2010) and taxonomy —in the latter through recognition of major contributions of revisions/monographs, although their citations are usually accumulated slow- and gradually (Venu & Sanjappa 2011). Longer articles are likely cited earlier because they have more citable content than shorter ones and may also have more visibility (Leimu & Koricheva 2005). For instance, during taxonomic revisions an entire genus—or family—can be revised, leading to discoveries of several new taxa or nomenclatural rearrangements (e.g., 47) that are subsequently cited by scientists working in the revised groups. However, this trend may not hold, or even be inverted, for papers published in high-impact multidisciplinary journals, such as *Science* and *Nature*, where most publications are usually only few-pages long. Additionally, ‘supplementary material’ may also contribute to the reduction of time to first citation since it provides additional and citable content attached to the main paper. The use of supplementary material is common ground in ecological works nowadays, but it is likely less so in taxonomy. Incorporating supplementary information into paper length would likely increase effect sizes.

Although we did not find significant differences between studies focused on different major taxonomic groups, the time to first citation varied greatly among groups (Figure 2a-b). Multitaxa studies had much lower times to first citation, particularly compared to studies focused on less conspicuous and charismatic organisms, where the overlap of survival curves and confidence intervals was minimal (Figure 2a). These studies may reach a larger audience as they ‘cover two different worlds’, that is, scientists working on very different taxonomic groups can potentially cite them. We acknowledge that our choice of grouping some organisms, but particularly, of using a high hierarchical level during analysis could affect our findings. For instance, a greater research effort to certain groups over others (Titley et al. 2017; Troudet et al. 2017) could ultimately influence first citation speed as well as other citation-based metrics. Future studies on this topic could use smaller taxonomic scales or even accommodate taxon identify as random structure into models. However, in the latter scenario researchers would not be interested in the particular effect of taxonomies upon the response variable under investigation, but only to control for dependence issues.

Contrarily to previous studies, we did not find an effect of collaboration on time to first citation across ecological and taxonomic articles. For instance, increased collaboration is associated with fast citation after publication in the fields of robotics and artificial intelligence (Kumari et al. 2020), music (Hancock 2015), and chemistry (Bornmann & Daniel 2010b). Multiauthorship likely contributes to wider dissemination of published articles within the scientific community and speeds up first citation (Bornmann & Daniel 2010b), but it may also be related to increased self-citations (Herbertz 1995; Leimu & Koricheva 2005). Differently from other studies, we removed self-citations herein, which likely contributed to the non-significance of the number of authors and countries on first citation speed. The impact of self-citations may be exacerbated across taxonomic articles, where it can play an important role in citation counts, particularly in more restricted research groups—e.g., an expert taxonomist working with an understudied and ‘understaffed’ group will likely end up citing himself/herself frequently.

## Conclusion

A plethora of scientometric indicators have been developed, many of which are based primarily on the traditional total number of citations [for a review, see (De Rijcke et al. 2016)]. These metrics are increasingly becoming central components of evaluation systems of scientific research and, consequently, of researchers themselves (Aksnes et al. 2019). In this study, we used the time to first citation to empirically demonstrate intrinsic differences between ecology and taxonomy. Despite their many differences, these two areas are directly related to each other (Freeman & Pennell 2021) and understating how scientometric indices behave in each of them may contribute, for instance, to a fairer distribution of research grants across disciplines. Despite the growing use and popularity of scientometric indicators, it is important to recognize which factors affect them, as well as their biases and pitfalls. Among future directions for studies on the time to first citation, we highlight the importance to investigate i) how different time intervals—from publication to cut-off dates—affects citation speed variation, ii) how the inclusion/exclusion of self-citations changes first citation times, particularly between different study areas, iii) how social and non-social media activity and related altimetrics affect citation speed, and iv) how article accessibility—open access *vs*. non-open access—is related to the time to first citation. Finally, we stress that no metric is perfect and the blind use of single traditional metrics to compare studies, researchers, or institutions can be highly pervasive. The combined use of several—and particularly unrelated—metrics is strongly recommended, especially for important decisions as grant allocation and career progression.

## Appendix

**Appendix 1**. List of ecological and taxonomic journals analyzed in this study. We selected 30 papers published during 2015 from each journal. *Ecology*: Acta Oecologica, Austral Ecology, Basic and Applied Ecology, Ecography, Ecology, Ecology Letters, Ecosphere, Journal of Applied Ecology, Oecologia, Oikos. *Taxonomy*: Brittonia, European journal of taxonomy, Mycotaxon, Phytotaxa, Systematic Botany, Taxon, Vertebrate Zoology, Zookeys, Zoological Journal of the Linnean Society, Zoosystema.

## Acknowledgments

We appreciate valuable friendly reviews in a previous version of this manuscript by Maurício Bini, João Nabout, José A. F. Diniz-Filho, and four anonymous reviewers from the DPAC course. We thank the Brazilian Coordenação de Aperfeiçoamento de Pessoal de Nível Superior (CAPES) and the Conselho Nacional de Desenvolvimento Científico e Tecnológico (CNPq) for financial support.

## Supplementary Information

**Figure S1:**
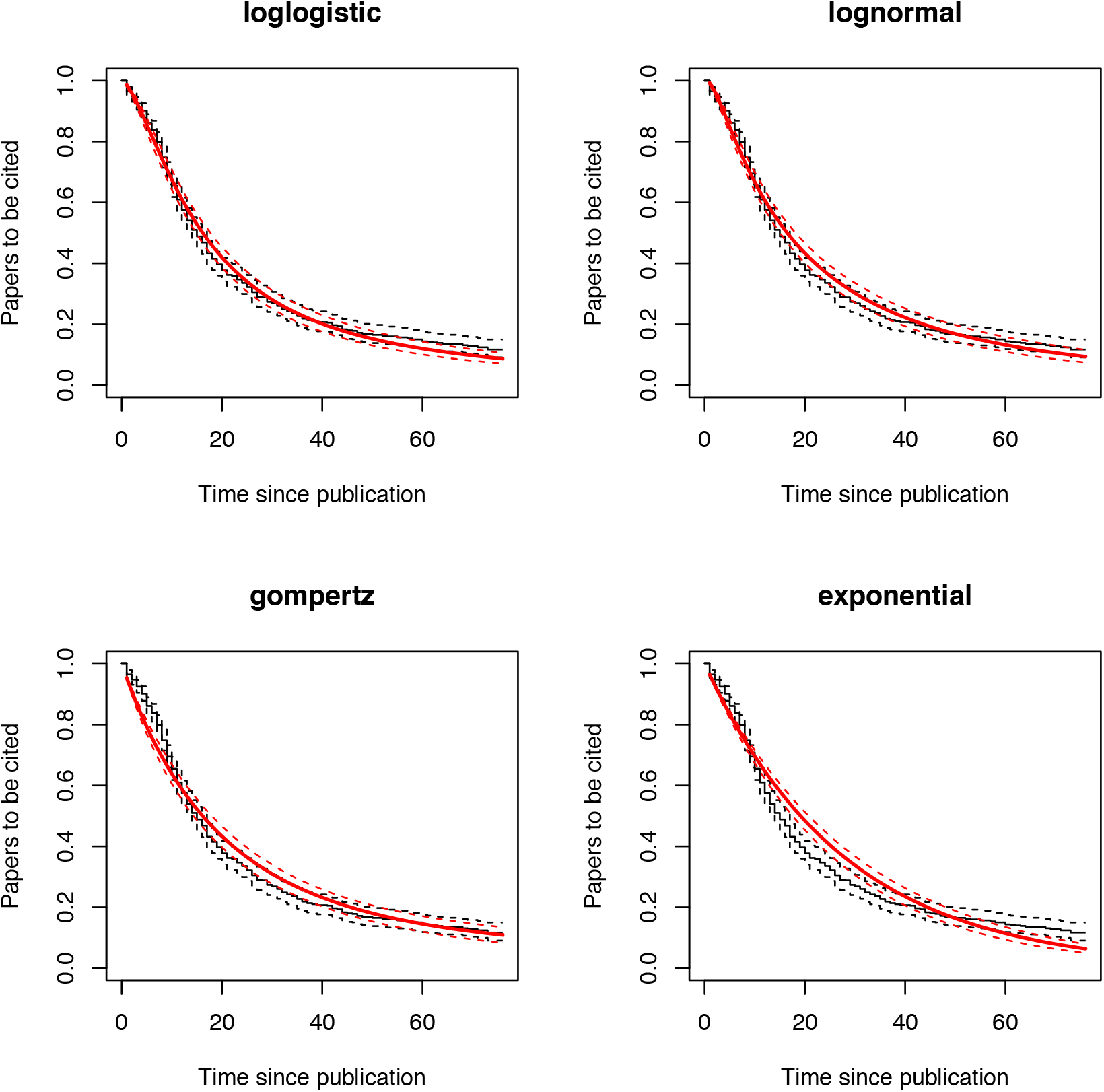
Top-4 best fitted family error distribution to the Accelerated Failure Time (AFT) survival model for the time to first citation in ecology and taxonomy. The best-fitted error distribution was the log logistic. See table S2 (below) for model parameters and AICc values.

**Figure S2:**
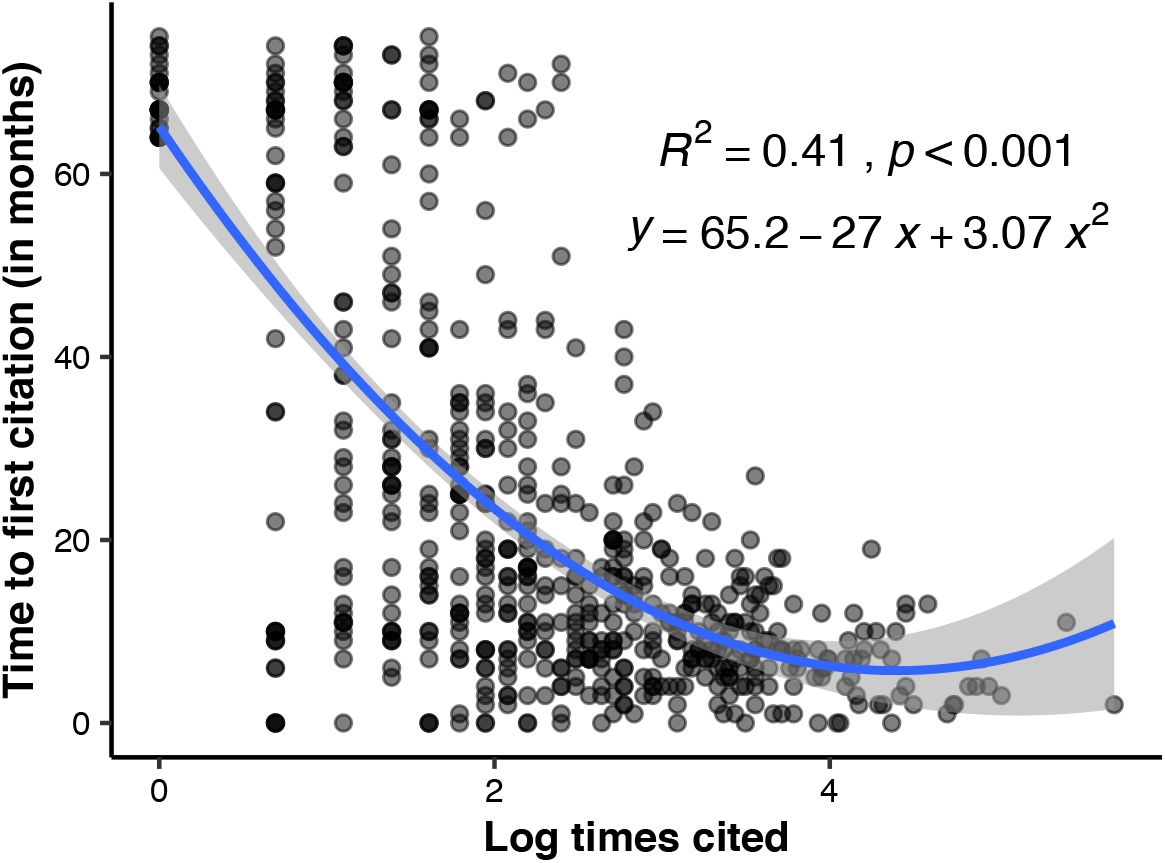
Correlation between the total number of citations (log-transformed) and the time to first citation across 600 ecological and taxonomic articles published in 2015. Adjusted correlation (in blue) was fitted using a second-order polynomial function and shows a negative relationship between the two metrics.

**Figure S3:**
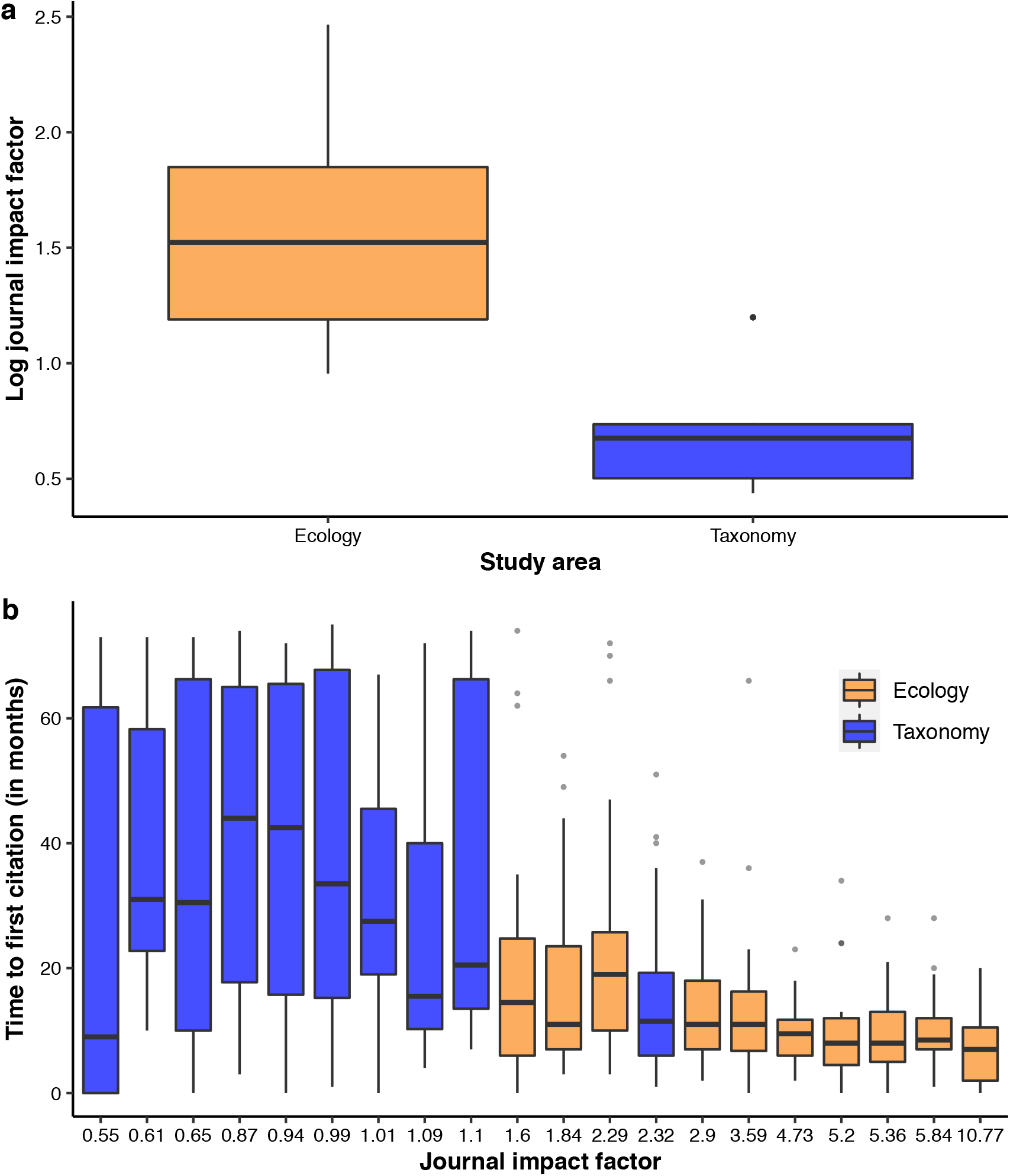
a) Differences in journal’s impact factors by area. b) The relationship between impact factor and time to first citation.

**Table S1:** Variation Inflation Factor among continuous predictors for the time to first citation in ecology and taxonomy.

| Variables | VIF |
| --- | --- |
| LogN.Pages.z | 1.040723 |
| LogN.Authors.z | 1.74842 |
| LogH.index.Max.z | 1.622732 |
| LogN.Countries.z | 1.468706 |
| GDP.Max.z | 1.367518 |

**Table S2:** Results from the model selection approach to identify the best family error distribution for the Accelerated Failure Time model. npars = are number of parameters for each distribution; AICc = Akaike information criteria; delta AICc = differences from best model; wAICc = model weights.

| **Distribution** | **npars** | **AICc** | **deltaAICc** | **wAICc** |
| --- | --- | --- | --- | --- |
| loglogistic | 2 | 4442.487 | 0.000 | 0.996 |
| lognormal | 2 | 4453.464 | 10.977 | 0.004 |
| gompertz | 2 | 4496.545 | 54.058 | 0.000 |
| exponential | 1 | 4527.628 | 85.141 | 0.000 |
| gamma | 2 | 4528.765 | 86.278 | 0.000 |
| weibull | 2 | 4528.978 | 86.491 | 0.000 |

